# Gonads or body? Differences in gonadal and somatic photoperiodic growth response in two vole species

**DOI:** 10.1101/2020.06.12.147777

**Authors:** Laura van Rosmalen, Jayme van Dalum, David G. Hazlerigg, Roelof A. Hut

## Abstract

To optimally time reproduction, seasonal mammals use a photoperiodic neuroendocrine system (PNES) that measures photoperiod and subsequently drives reproduction. To adapt to late spring arrival at northern latitudes, a lower photoperiodic sensitivity and therefore a higher critical photoperiod for reproductive onset is necessary in northern species to arrest reproductive development until spring onset. Temperature-photoperiod relationships, and hence food availability-photoperiod relationships, are highly latitude dependent. Therefore, we predict PNES sensitivity characteristics to be latitude-dependent. Here, we investigated photoperiodic responses at different times during development in northern- (tundra/root vole, *Microtus oeconomus*) and southern vole species (common vole, *Microtus arvalis*) exposed to constant short (SP) or long photoperiod (LP).

*M. oeconomus* grows faster under LP, whereas no photoperiodic effect on somatic growth is observed in *M. arvalis*. Contrastingly, gonadal growth is more sensitive to photoperiod in *M. arvalis*, suggesting that photoperiodic responses in somatic and gonadal growth can be plastic, and might be regulated through different mechanisms. In both species, thyroid-stimulating-hormone-β subunit (*Tshβ*) and iodothyronine-deiodinase 2 (*Dio2*) expression is highly increased under LP, whereas *Tshr* and *Dio3* decreases under LP. High *Tshr* levels in voles raised under SP may lead to increased sensitivity to increasing photoperiods later in life. The higher photoperiodic induced *Tshr* response in *M. oeconomus* suggests that the northern vole species might be more sensitive to TSH when raised under SP.

Species differences in developmental programming of the PNES, which is dependent on photoperiod early in development, may form part divergent breeding strategies evolving as part of latitudinal adaptation.

**Summary statement:** Development of the neuroendocrine system driving photoperiodic responses in gonadal and somatic growth differ between the common and the tundra vole, indicating that they use a different breeding strategy.

## Introduction

Organisms use intrinsic annual timing mechanisms to adaptively prepare behavior, physiology, and morphology for the upcoming season. In temperate regions, decreased ambient temperature is associated with reduced food availability during winter which will impose increased energetic challenges that completely prevent the possibility of successfully raising offspring. Annual variation in ambient temperature shows large fluctuations between years, with considerable day to day variations, whereas annual changes in photoperiod provide a consistent year-on-year signal for annual phase. This has led to convergent evolutionary processes in many organisms to use day length as the most reliable cue for seasonal adaptations.

In mammals, the photoperiodic neuroendocrine system (PNES) measures photoperiod and subsequently drives annual rhythms in physiology and reproduction (for review see Dardente *et al*., 2018; Nakane and Yoshimura, 2019). Light is perceived by retinal photoreceptors that signal to the suprachiasmatic nucleus (SCN). The SCN acts on the pineal gland, such that the duration of melatonin production during darkness changes over the year to represent the inverse of day length. Melatonin binds to its receptor (MTNR1A) in the pars tuberalis (PT) of the anterior lobe of the pituitary gland (Gall et al., 2002; Gall et al., 2005; Williams and Morgan, 1988). Under long days, pineal melatonin is released for a short duration and thyroid stimulating hormone β subunit (*Tshβ)* expression is increased in the pars tuberalis, leading to increased secretion of TSH. PT-derived TSH acts locally through TSH receptors (TSHr) found in the tanycytes in the neighbouring mediobasal hypothalamus (MBH). The tanycytes produce increased iodothyronine deiodinase 2 (DIO2) and decreased DIO3 levels, which leads to higher levels of the active form of thyroid hormone (T_3_) and lower levels of inactive forms of thyroid hormone (T4 and rT3). In small mammals, it is likely that T_3_ acts ‘indirectly’, through KNDy (kisspeptin/neurokininB/Dynorphin) neurons of the arcuate nucleus (ARC) (for review see Simonneaux, 2020) in turn controlling the activity of gonadotropin-releasing hormone (GnRH) neurons. GnRH neurons project to the pituitary to induce gonadotropin release, which stimulates gonadal growth. The neuroanatomy of this mechanism has been mapped in detail and genes and promoter elements that play a crucial role in this response pathway have been identified in several mammalian species (Dardente et al., 2010; Hanon et al., 2008; Hut, 2011; Masumoto et al., 2010; Nakao et al., 2008; Ono et al., 2008; Sáenz De Miera et al., 2014; Wood et al., 2015), including the common vole (Król et al., 2012).

Voles are small grass-eating rodents with a short gestation time (i.e. 21 days). They can have several litters a year, while their offspring can reach sexual maturity within a month during spring and summer. Depending on time of year, pups need to either accelerate reproductive development in spring and summer, or to delay reproductive development in late summer and autumn. In small rodents, photoperiods experienced in utero, already prior to birth, determines growth rate and reproductive development. Photoperiodic reactions to intermediate day lengths depend on prior photoperiodic exposure during both prenatal and postnatal life stages (Hoffmann, 1973; Horton, 1984a; Horton, 1984b; Horton, 1985; Horton and Stetson, 1992; Prendergast et al., 2000; Sáenz de Miera et al., 2017; Stetson et al., 1986; Yellon and Goldman, 1984). Maternal photoperiodic information is transferred to the young in utero by melatonin in several rodent species (Horton and Stetson, 1992; Yellon and Longo, 1987). By passing information about day length from mother to fetus, offspring will be prepared for the upcoming season. Presumably, crucial photoperiod-dependent steps in PNES development take place in young animals to secure an appropriate seasonal response later in life (Dalum et al., 2020; Sáenz de Miera et al., 2017; Sáenz de Miera et al., 2020; Sáenz De Miera, 2019). In Siberian hamsters, maternal programming takes place downstream of melatonin secretion at the level of *Tshr*, with expression increased in animals born under SP, associated with subsequent increases in TSH sensitivity (Sáenz de Miera et al., 2017).

Because primary production in the food web of terrestrial ecosystems is temperature-dependent (Robson, 1967; Peacock, 1976; Malyshev *et al*., 2014), we can expect herbivores like *Microtus* species to adjust their photoperiodic response such that reproduction in spring starts when primary food production starts (Baker, 1938). *Microtus* is a genus of voles found in the northern hemisphere, ranging from close to the equator to arctic regions, which makes it an excellent species to study latitudinal adaptation to photoperiodic responses. Photoperiodically induced reproduction should start at longer photoperiods at more northern populations, since a specific ambient spring temperature at higher latitudes coincides with longer photoperiods compared to lower latitudes (Hut et al., 2013). To adapt to late spring arrival at northern latitudes, a lower sensitivity to photoperiod and therefore a longer critical photoperiod is necessary in northern species to arrest reproductive development until spring onset. (epi)genetic adaptation to local annual environmental changes may create latitudinal differences in photoperiodic responses and annual timing mechanisms.

Populations with different latitudinal distributions, such as *Microtus* populations, can be useful to understand adaptation of photoperiodic responses (for review see Hut et al., 2013). In order to understand the development of the PNES for vole species with different paleogeographic origins, we investigated photoperiodic responses at different time points during development by exposing northern- (tundra/root vole, *Microtus oeconomus*) and southern vole species (common vole, *Microtus* arvalis) to constant short- or long photoperiods in the laboratory. Animals from our two vole lab populations are originally caught from the same latitude in the Netherlands (53°N) where both populations overlap. This is for *M. arvalis* the center (mid-latitude) of its distribution range (38-62°N), while our lab *M. oeconomus* are from a relict population at the lower boundary of its European geographical range (48-72°N). This leads to the expectation that the PNES of *M. arvalis* is better adapted to the local annual environmental changes of the original latitude, whereas *M. oeconomus* might show a photoperiodic response characteristic of more northern regions.

## Materials and methods

### Animals and experimental procedures

All experimental procedures were carried out according to the guidelines of the animal welfare body (IvD) of the University of Groningen, and all experiments were approved by the Centrale Commissie Dierproeven) of the Netherlands (CCD, license number: AVD1050020171566). The Groningen common vole breeding colony started with voles (*M. arvalis* (Pallas, 1778)) obtained from the Lauwersmeer area (Netherlands, 53° 24’ N, 6° 16’ E) (Gerkema et al., 1993), and was occasionally supplemented with wild caught voles from the same region to prevent the lab population from inbreeding. The Groningen tundra vole colony started with voles (*M. oeconomus* (Pallas, 1776)) obtained from four different regions in the Netherlands (described in Van de Zande et al., 2000). Both breeding colonies were maintained at the University of Groningen as outbred colonies and provided the voles for this study. All breeding pairs were kept in climate controlled rooms, at an ambient temperature of 21 ±1°C and 55 ±5% relative humidity and housed in transparent plastic cages (15 x 40 x 24 cm) provided with sawdust, dried hay, an opaque pvc tube and *ad libitum* water and food (standard rodent chow, #141005; Altromin International, Lage, Germany).

The voles used in the experiments (61 males, 56 females) were both gestated and born under either a long photoperiod (LP, 16h light: 8h dark) or a short photoperiod (SP, 8h light: 16h dark). Pups were weaned and transferred to individual cages (15 x 40 x 24 cm) when 21 days old but remained exposed to the same photoperiod as during both gestation and birth. All voles were weighed at post-natal day 7, 15, 21, 30, 42 and 50 (Fig. 1).

**Figure 1.**
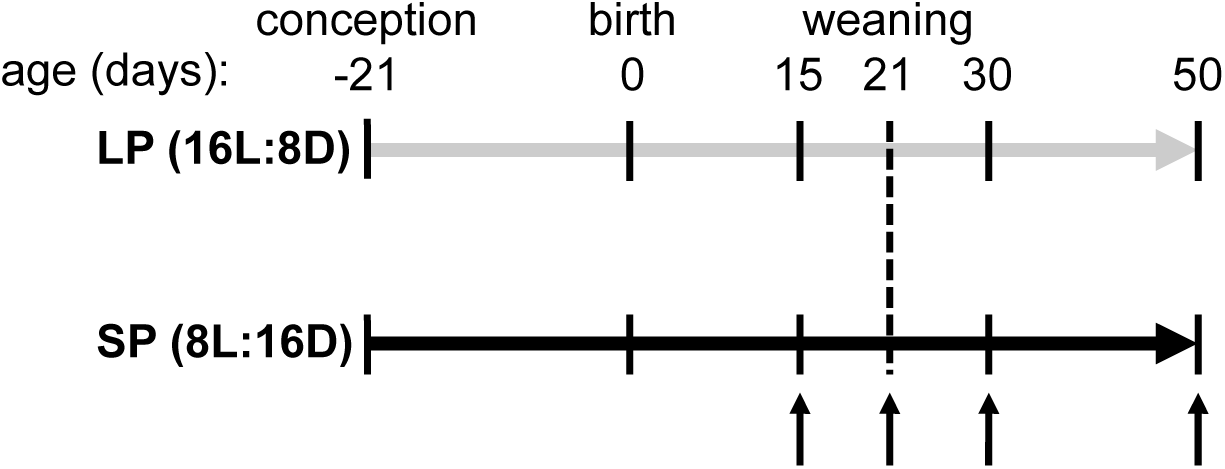
Experimental design. Animals were constantly exposed to either LP or SP from gestation onwards. Arrows indicate sampling points for tissue collection. Age in days is depicted above the timeline. Vertical dashed line represents time of weaning (21 days old).

### Tissue collections

In order to follow development, animals were sacrificed by decapitation 16±1 hours after lights off (*Tshβ* expression peaking in pars tuberalis (Masumoto et al., 2010)), at an age of 15, 21, 30 and 50 days old. Brains were removed with great care to include stalk of the pituitary containing the pars tuberalis. The hypothalamus with the pars tuberalis were dissected as described in Prendergast et al., 2013: the optic chiasm at the anterior border, the mammillary bodies at the posterior border, and laterally at the hypothalamic sulci. The remaining hypothalamic block was cut dorsally 3-4 mm from the ventral surface. The extracted hypothalamic tissue was flash frozen in liquid N_2_ and stored at -80°C until RNA extraction. Reproductive organs were dissected and cleaned of fat, and wet masses of paired testis, paired ovary and uterus were measured (±0.0001 g).

### RNA extraction, Reverse Transcription and Real-time quantitative PCR

Total RNA was isolated from the dissected part of the hypothalamus using TRIzol reagent according to the manufacturer’s protocol (Invitrogen™, Carlsbad, California, United States). In short, frozen pieces of tissue (∼0.02 g) were homogenized in 0.5 ml TRIzol reagent in a TissueLyser II (Qiagen, Hilden, Germany) (2 x 2 minutes at 30 Hz) using tubes containing a 5mm RNase free stainless-steel bead. Subsequently 0.1 ml chloroform was added for phase separation. Following RNA precipitation by 0.25 ml of 100% isopropanol, the obtained pellet was washed with 0.5 ml of 75% ETOH. Depending on the size, RNA pellets were diluted in an adequate volume of RNase-free H_2_O (range 20-50 µL) and quantified on a Nanodrop 2000 (Thermoscientific™, Waltham, Massachusetts, United States). After DNA removal by DNase I treatment (Invitrogen™, Carlsbad, California, United States), equal quantity of RNA from each sample was used for cDNA synthesis by using RevertAid H minus first strand cDNA synthesis reagents (Thermoscientific™, Waltham, Massachusetts, United States). 40 µL Reverse Transcription (RT) reactions were prepared using 2 µg RNA, 100 µM Oligo(dT)^18^, 5x Reaction buffer, 20 U/µL RiboLock RNase Inhibitor, 10 mM dNTP Mix, RevertAid H Minus Reverse Transcriptase (200 U/µL). Concentrations used for RT reactions can be found in the supplementary information (table S1). RNA was reversed transcribed by using a thermal cycler (S1000™, Bio-Rad, Hercules, California, United States). Incubation conditions used for RT were: 45°C for 60 minutes followed by 70°C for 5 minutes. Transcript levels were quantified by Real-Time qPCR using SYBR Green (KAPA SYBR FAST qPCR Master Mix, Kapa Biosystems). 20 μL (2 μL cDNA + 18 μL Mastermix) reactions were carried out in duplo for each sample by using 96-well plates in a Fast Real-Time PCR System (CFX96, Bio-Rad, Hercules, California, United States). Primers for genes of interest were designed using Primer-BLAST (NCBI) and optimized annealing temperature (Tm) and primer concentration. All primers used in this study are listed in the supplementary information (table S2). Thermal cycling conditions used can be found in the supplementary information (table S3). Relative mRNA expression levels were calculated based on the ΔΔCT method using *Gapdh* as the reference (housekeeping) gene (Pfaffl 2001).

### Statistical analysis

Sample size (n = 4) was determined by a power calculation (α = 0.05, power = 0.80) based on the effect size (d = 2.53) of an earlier study, in which gonadal weight was assessed in voles under three different photoperiods (Król et al., 2012). Effects of age, photoperiod and species on body mass, reproductive organs and gene expression levels were determined using a two-way ANOVA. Tukey HSD post-hoc pairwise comparisons were used to compare groups at specific ages. Statistical significance was determined at *p* < 0.05. Statistical results can be found in the supplementary information (table S4). All statistical analyses were performed using RStudio (version 1.2.1335), and figures were generated using the ggplot2 package.

## Results

### Body mass responses for males and females

Photoperiod during gestation did not affect birth weight in either species (Fig. 2A,B). Both *M. oeconomus* males and females grow faster under LP compared to SP conditions (males, *F*_1,303_ = 15.0, *p* < 0.001; females, *F*_1,307_ = 10.2, *p* < 0.01) (Fig. 2A,B). However, no effect of photoperiod on body mass was observed in *M. arvalis* males or females (males, *F*_1,243_ = 2.1, ns; females, *F*_1,234_ = 0.6, ns) (Fig. 2A,B).

**Figure 2.**
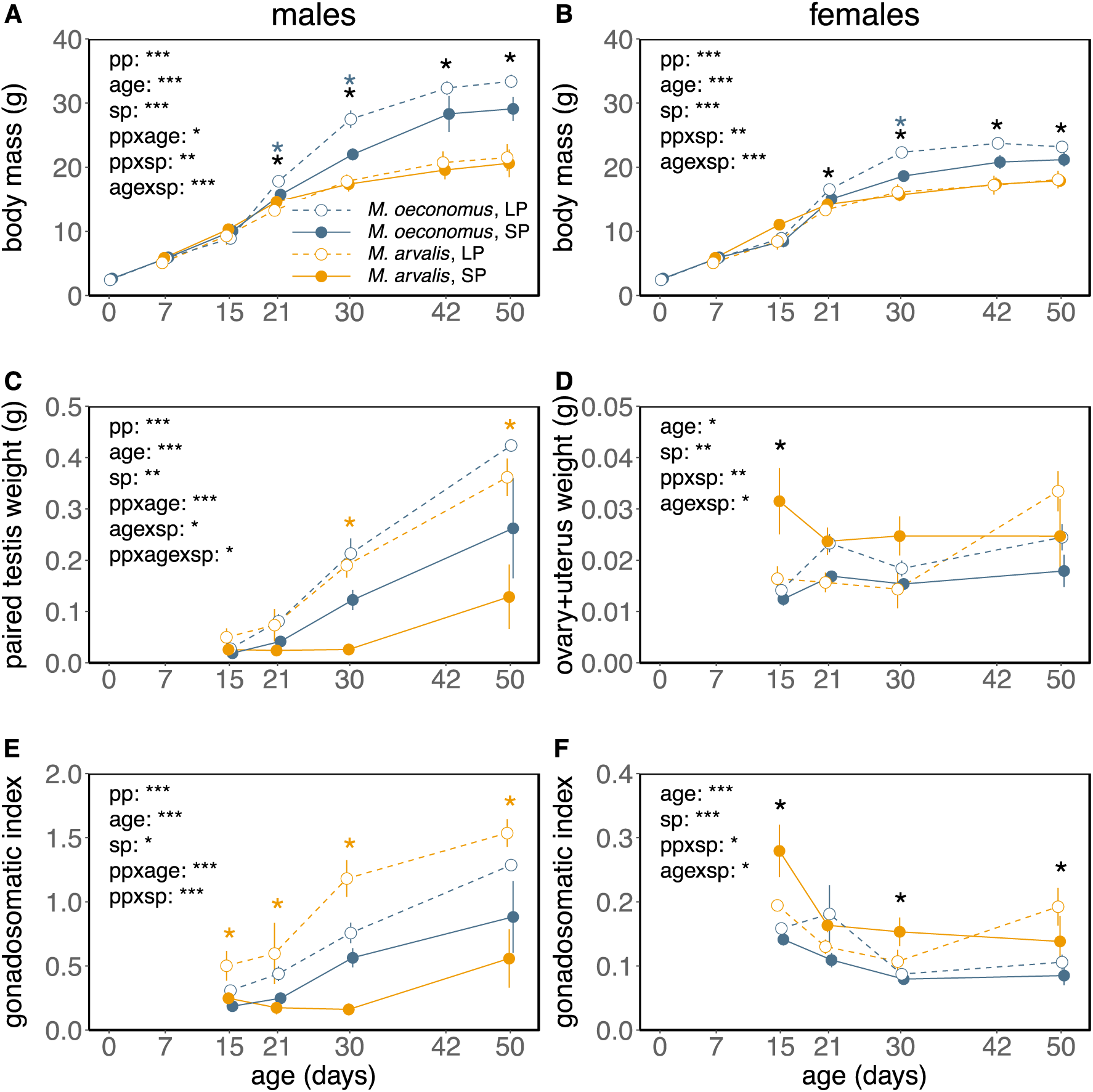
Effects of constant photoperiod on growth and gonadal development. Graphs depicting body mass growth curves for (A) males and (B) females, (C) paired testis weight, (D) paired ovary + uterus weight, (E, F) gonadal development relative to body mass (gonadosomatic index) for *M. arvalis* (orange) and *M. oeconomus* (blue), continuously exposed to either LP (open symbols, dashed lines) or SP (closed symbols, solid lines). Data are mean±s.e.m. Significant effects (two-way ANOVA’s, post-hoc Tukey) of photoperiod at specific ages are indicated for *M. oeconomus* (blue asterisks) and *M. arvalis* (orange asterisks), significant effects of species are indicated by black asterisks. Significant effects of: photoperiod (pp), age (age), species (sp) and interactions are shown in each graph, *p < 0.05, **p < 0.01, ***p < 0.001. Statistic results for two-way ANOVA’s (photoperiod, age and species) can be found in table S4.

### Gonadal responses for males

*M. arvalis* males show faster testis growth under LP compared to SP (testis, *F*_1,33_ = 17.01, *p* < 0.001; GSI, *F*_1,33_ = 32.2, *p* < 0.001) (Fig. 2C,E). This photoperiodic effect on testis development is less pronounced in *M. oeconomus* (testis, *F*_1,35_ = 8.3, *p* < 0.01; GSI, *F*_1,35_ = 9.3, *p* < 0.01) (Fig. 2C,E).

### Gonadal responses for females

*M. arvalis* female gonadal weight (i.e. paired ovary + uterus) is slightly higher in the beginning of development (until 30 days old) under SP compared to LP conditions (*F*_1,17_ = 10.4, *p* < 0.01) (Fig. 2D), while the opposite effect was observed in *M. oeconomus* (*F*_1,36_ = 9.0, *p* < 0.01) (Fig. 2D). For both species, these photoperiodic effects disappeared when gonadal mass was corrected for body mass (*M. arvalis, F*_1,17_ = 2.5, ns; *M. oeconomus, F*_1,36_ = 2.3, ns) (Fig. 2F). Interestingly, gonadal weight is significantly increasing in 30-50 days old LP *M. arvalis* females (*F*_1,5_ = 7.7, *p* < 0.05) (Fig. 2D), but not in *M. oeconomus* (*F*_1,11_ = 2.2, ns) or under SP conditions (*M. arvalis, F*_1,7_ = 0, ns; *M. oeconomus, F*_1,7_ = 1.0, ns).

### Photoperiod induced changes in hypothalamic gene expression

In males of both species, *Mtnr1a* expression in the hypothalamic block with preserved pars tuberalis was highly expressed, but unaffected by photoperiod or age (photoperiod, *F*_1,43_ = 0.08, ns; age, *F*_3,42_ = 0.94, ns) (Fig. 3A). In females, *Mtnr1a* expression increases approximately 2-fold with age in both species (*F*_3,40_ = 9.04, *p* < 0.001) (Fig. 3B), but no effects of photoperiod where observed (*F*_1,40_ =1.59, ns).

**Figure 3.**
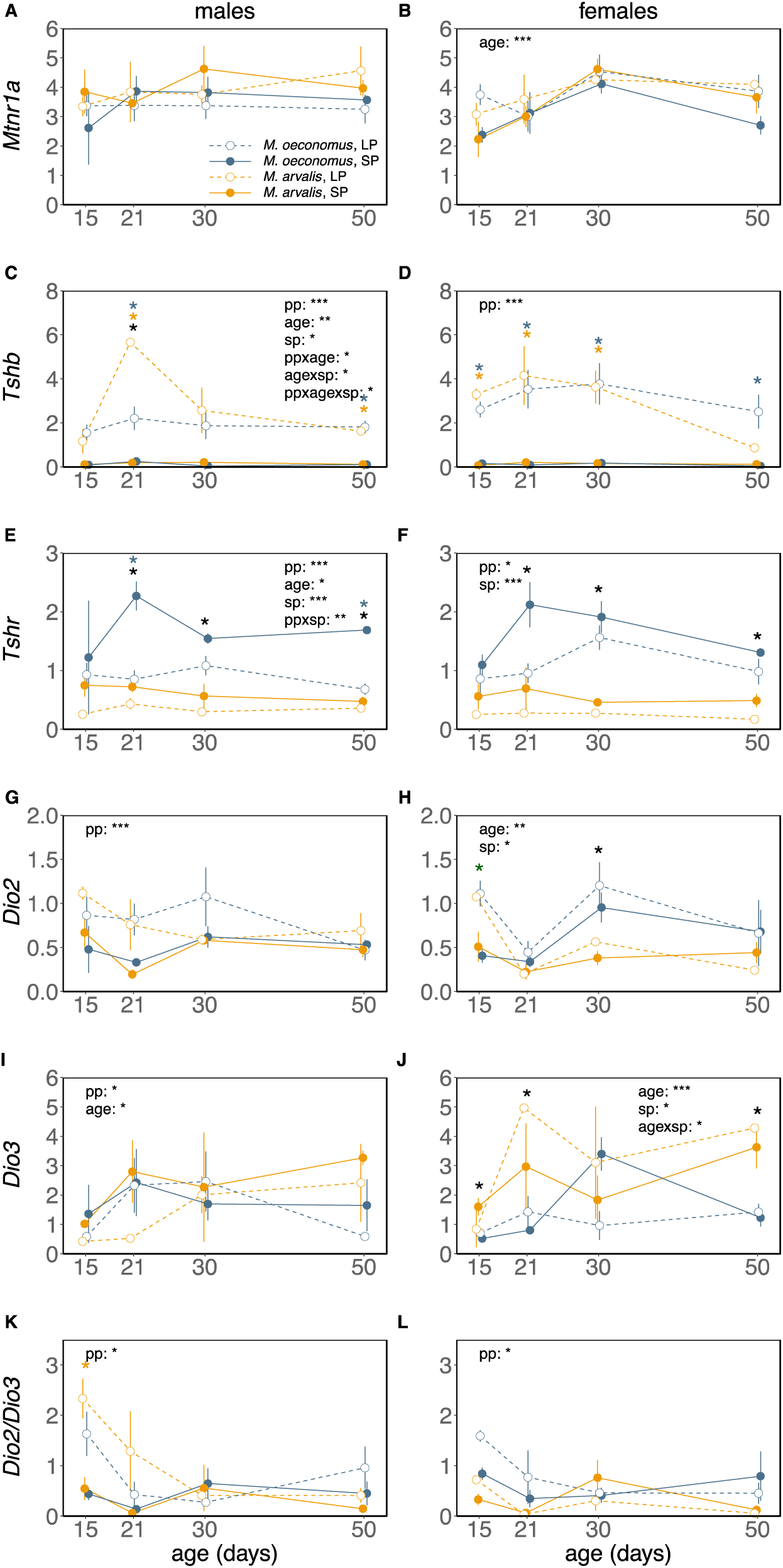
Effects of constant photoperiod on gene expression levels in the developing hypothalamus. Relative gene expression levels of (A, B) *Mtnr1a*, (C, D) *Tshβ*, (E, F) *Tshr*, (G, H) *Dio2*, (I, J) *Dio3* and (K, L) *Dio2/Dio3* expression in the hypothalamus of developing *M. arvalis* (orange) and *M. oeconomus* (blue) males and females respectively, under LP (open symbols, dashed lines) or SP (closed symbols, solid lines). Data are mean±s.e.m (n = 2-5). Significant effects (two-way ANOVA’s, post-hoc Tukey) of photoperiod at specific ages are indicate for *M. oeconomus* (blue asterisks) and *M. arvalis* (orange asterisks), significant effects of species are indicated by black asterisks. Significant effects of: photoperiod (pp), age (age), species (sp) and interactions are shown in each graph, *p < 0.05, **p < 0.01, ***p < 0.001. Statistic results for two way ANOVA’s (photoperiod, age and species) can be found in table S4.

In males and females of both species, *Tshβ* expression is dramatically elevated under LP throughout development (*M. oeconomus* males, *F*_1,27_ = 49.3, *p* < 0.001; *M. arvalis* males, *F*_1,27_ = 21.3, *p* < 0.001; *M. oeconomus* females, *F*_1,30_ = 63.7, *p* < 0.001; *M. arvalis* females, *F*_1,22_ = 60.9, *p* < 0.001) (Fig. 3C, D). Furthermore, a clear peak in *Tshβ* expression is observed in 21-day old LP *M. arvalis* males, while such a peak is lacking in *M. oeconomus* males. On the other hand, *Tshβ* expression in *M. oeconomus* males remains similar over the course of development under LP conditions. In females, photoperiodic responses on *Tshβ* expression did not differ between species (*F*_1,40_ = 0.02, ns).

In *M. oeconomus* males and females, *Tshr* expression is higher under SP compared to LP (males, *F*_1,27_ = 23.7, *p* < 0.001; females, *F*_1,30_ = 6.2, *p* < 0.05) (Fig. 3E,F), while photoperiodic induced changes in *Tshr* expression are smaller in *M. arvalis* males and females (males, *F*_1,27_ = 23.7, *p* < 0.01; females, *F*_1,22_ = 4.3, *p* < 0.05) (Fig. 3E,F). Photoperiodic responses on *Tshr* expression are significantly larger in *M. oeconomus* males compared to *M. arvalis* males (*F*_1,42_ = 8.17, *p* < 0.01) (Fig. 3E).

In males of both species, the largest photoperiodic effect on *Dio2* is found at weaning (day 21), with higher levels under LP compared to SP (*F*_1,42_ = 14.7, *p* < 0.001) (Fig. 3G). Interestingly, *Dio3* is lower in these animals (*F*_1,42_ = 4.8, *p* < 0.05) (Fig. 3I), leading to a high *Dio2/Dio3* ratio under LP in the beginning of development (*F*_1,42_ = 8.5, *p* < 0.01) (Fig. 3K). We find a similar pattern in females, with higher *Dio2* under LP compared to SP at the beginning of development (i.e. day 15) (*F*_3,10_ = 8.9, *p* < 0.01) (Fig. 3H).

In males of both species, no effects of photoperiod on Eyes Absent 3 (*Eya3*) (*F*_1,42_ = 1.72, ns), Kisspeptin (*Kiss1*) (*F*_1,42_ = 2.96, ns), Neuropeptide VF precursor (*Npvf*) (*F*_1,42_ = 0.61, ns), DNA methyltransferase 1 (*Dnmt1*) (*F*_1,42_ = 0.68, ns) and *Dnmt3a* (*F*_1,42_ = 0.77, ns) expression were found (Fig. S1A-I). In females, both *Kiss1* (*F*_3,40_ = 4.82, *p* < 0.01) and *Npvf* is higher under LP dependent on age (*F*_3,40_ = 3.51, *p* < 0.05) (Fig. S1D,F), but there were no effects of photoperiod on *Eya3* (*F*_1,40_ = 0.30, ns), *Dntm1* (*F*_1,40_ = 0.18, ns) and *Dnmt3a* (*F*_1,40_ = 0.08, ns) (Fig. S1B,H,J).

A positive correlation between the levels of *Tshβ* and *Dio2* expression was found only at the beginning of development (15 days, *F*_1,25_ = 12.6, *p* < 0.01; 21 days, *F*_1,28_ = 4.0, *p* < 0.1; 30 days, *F*_1,30_ = 0.1, ns; 50 days, *F*_1,23_ = 0.1, ns) (Fig. 4A-D). Moreover, no significant relationship between *Dio2* and *Dio3* expression was found (Fig. 4E-H).

**Figure 4.**
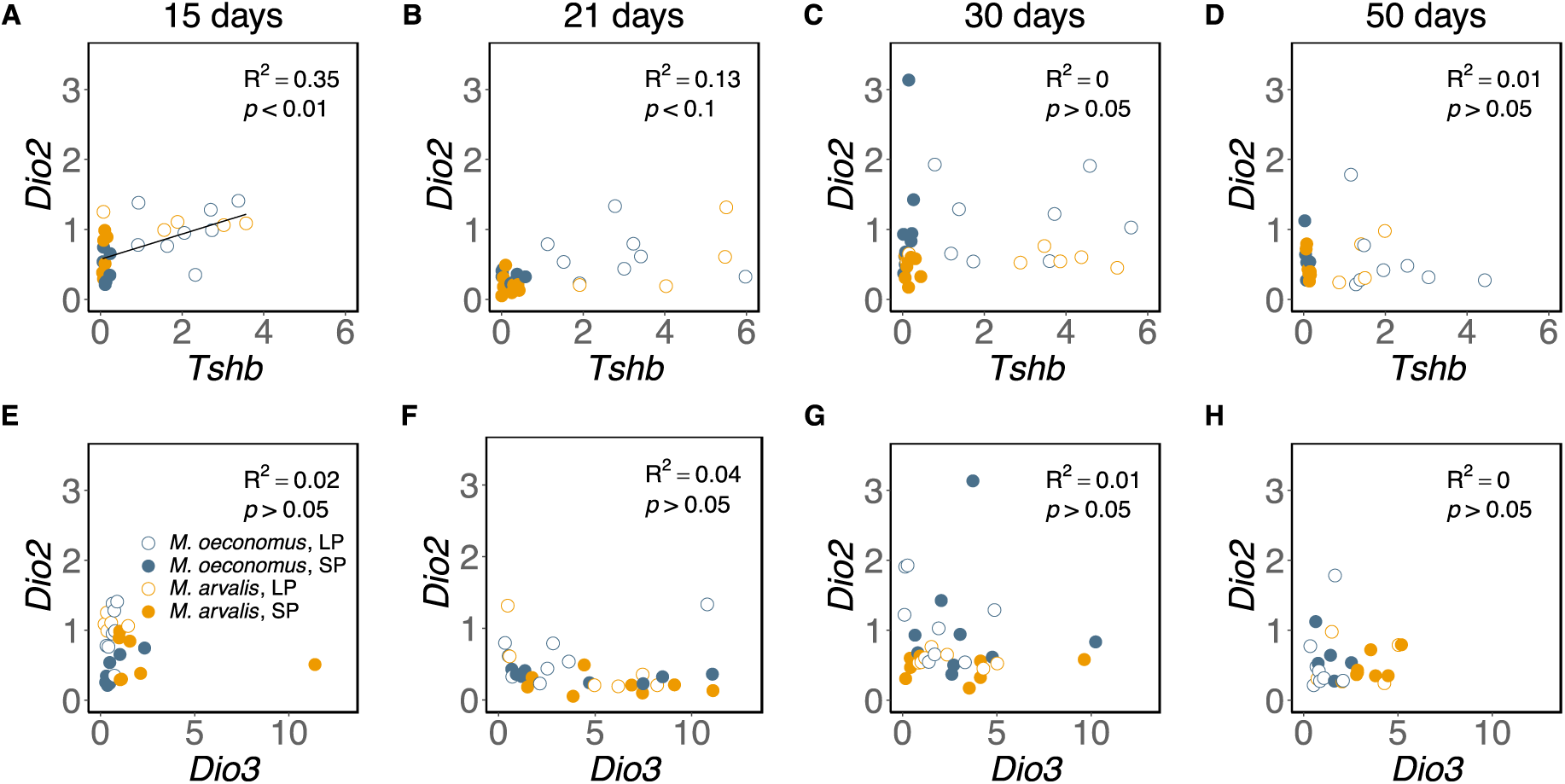
Relationship between hypothalamic *Dio2, Dio3* and *Tshβ* expression in voles at different age. Scatterplot of *Tshβ* versus *Dio2* gene expression at (A) 15, (B) 21, (C) 30 and (D) 50 days old. Scatterplot of *Dio3* versus *Dio2* gene expression at (E) 15, (F) 21, (G) 30 and (H) 50 days old. Open symbols indicate LP animals, closed symbols indicate SP animals. blue symbols represent *M. oeconomus*, orange symbols represent *M. arvalis*.

## Discussion

This study demonstrates different effects of constant photoperiod on the PNES in two different *Microtus* species: *M. arvalis* and *M. oeconomus*. Overall, somatic growth is photoperiodically sensitive in *M. oeconomus* while gonadal growth is photoperiodically sensitive in *M. arvalis*. Hypothalamic *Tshβ, Tshr, Dio2* and *Dio3* expression is highly affected by photoperiod and age, and some species differences were observed in the magnitude of these effects.

### Photoperiod induced changes in somatic growth and gonadal development

These data demonstrate that photoperiod early in life affects pup growth in *M. oeconomus* (Fig. 2A), and reproductive development in *M. arvalis* males (Fig. 2C,E). In females, a similar photoperiodic effect on somatic growth is observed as in males. *M. oeconomus* females grow faster under LP compared to SP, while there is no difference in growth rates between LP and SP in *M. arvalis* (Fig. 2B). In *M. oeconomus*, somatic growth is plastic, whereas, in *M. arvalis*, gonadal growth is plastic. Garden dormouse (*Eliomys quercinus*) born late in the season grow and fatten twice as fast as early born animals (Stumpfel et al., 2017), in order to partly compensate for the limited time before winter onset. This overwintering strategy might be favorable for animals with a short breeding season (i.e. high latitude), and may also be used in *M. oeconomus* since they gain weight faster when raised under LP (i.e. late in the season) compared to SP (i.e. early in the season). Southern arvicoline species have longer breeding seasons (Tkadlec, 2000), and therefore have more time left to compensate body mass when born late in de season. Therefore, somatic growth rate may depend to a lesser extent on the timing of birth in southern species as observed in *M. arvalis* raised under SP or LP.

*M. arvalis* female gonadal weight is slightly higher under SP compared to LP at the beginning of development (Fig. 2D,F). In contrast, hamster uterus weight is increased after 3 weeks of constant LP exposure, which continued throughout development (Ebling, 1994; Phalen et al., 2009). In *M. arvalis*, female gonadal weight is increasing from day 30 to day 50 in LP animals, whereas gonadal weight in SP females remains the same (Fig. 2D,F). Also, *M. oeconomus* female gonadal weight is not increased in this period of development under both LP and SP conditions. Puberty onset in *M. arvalis* is later in time compared to Siberian hamsters (Phalen et al., 2009), while earlier in time compared to *M. oeconomus*, and thus LP *M. arvalis* increases gonadal weight earlier in development (i.e. > 30 days old) compared to LP *M. oeconomus* (i.e. > 50 days old), in order to increase reproductive activity and prepare for pregnancy. These results suggest that *M. oeconomus* females have a later reproductive onset compared to *M. arvalis* females under LP conditions, indicating that a photoperiod of 16L:8D is detected as a spring photoperiod for *M. arvalis*, whereas a longer photoperiod may be needed to induce reproductive development in *M. oeconomus*. This hypothesis can be tested by exposing voles to a broader range of different photoperiod regimes in the laboratory. Our data shows that *M. arvalis* invest more energy into gonadal growth, whereas *M. oeconomus* invest more energy into body mass growth independent of gonadal growth under LP. This suggests that both body mass growth and gonadal development are plastic and can be differentially affected by photoperiod, perhaps through different mechanisms. In Siberian hamsters, the growth hormone (GH) axis is involved in photoperiodic regulation of body mass (Dumbell et al., 2015; Scherbarth et al., 2015). Our results indicate a different role for the GH-axis in seasonal body mass regulation in *M. oeconomus* and *M. arvalis*.

### Photoperiod induced changes in hypothalamic gene expression

*M. arvalis* males show a clear photoperiodic response in both hypothalamic gene expression and gonadal activation. Genes in the female PNES are strongly regulated by photoperiod, which is not reflected in gonadal growth. In *M. oeconomus*, PNES gene expression profiles change accordingly to photoperiod, however the gonadal response is less sensitive to photoperiod, which is similar to the photoperiodic response observed in house mice (Masumoto et al., 2010). Because *M. oeconomus* is more common at high latitudes, where they live in tunnels covered by snow in winter and early spring, photoperiodic information might be blocked during a large part of the year for these animals (Evernden and Fuller, 1972; Korslund, 2006). For this reason, other environmental cues, such as metabolic status, may integrate in the PNES in order to regulate the gonadal response and therefore timing of reproduction. Because, female voles do not show a strong gonadal response to photoperiod either, a similar mechanism as in *M. oeconomus* is expected.

### Photoperiod induced changes in Tshβ sensitivity

In both vole species *Tshβ* expression is higher under LP conditions during all stages of development (Fig. 3C,D), which is in agreement with previous studies in other mammals, birds and fish (for review see Dardente et al., 2014; Nakane and Yoshimura, 2019). We sampled 16 hours after lights off, when *Tshβ* expression is peaking. Perhaps we sampled too late in order to find photoperiodic induced changes in *Eya3* expression (Fig. S1A,B), since in mice *Eya3* is peaking 12 hours after lights off under LP conditions (Masumoto et al., 2010).

Although, less pronounced in *M. arvalis*, elevated *Tshr* expression under SP (Fig. 3E,F) may be caused by low *Tshβ* levels in the same animals (Fig. 3C,D). In a previous study, a similar relationship between *Tshr* and *Tshβ* expression in the pars tuberalis and medial basal hypothalamus (MBH) of Siberian hamsters has been observed (Sáenz de Miera et al., 2017). In our study, the ependymal paraventricular zone (PVZ) around the third ventricle of the brain and the pars tuberalis are both included in samples for RNA extraction and qPCR, therefore, we cannot distinguish between these two brain areas. Brains were collected 16 hours after lights off, when *Tshr* mRNA levels in the pars tuberalis and PVZ are predicted to be similar based on studies in sheep (Hanon et al., 2008). Similar circadian expression patterns are expected in brains of seasonal long-day breeding rodents. Therefore, the observed increase in *Tshr* expression in SP voles, of both species and sexes, (Fig. 3E,F) may relate to high TSH density in the tanycytes lining the third ventricle, which might lead to increased TSH sensitivity later in life. The high *Tshr* expression in voles developing under SP (Fig. 3E,F) may favour a heightened sensitivity to increasing TSH, photoperiods increase later in life. This in turn would promote increased DIO2 and decreased DIO3 levels in spring. Interestingly, photoperiodic responses on *Tshr* are more pronounced in *M. oeconomus* than in *M. arvalis*, suggesting that *M. oeconomus* is more sensitive to TSH protein when raised under SP. However, TSH is a dimer of αGSU and TSHβ, and we did not measure αGSU levels in this study.

Our vole lab populations are originally from the same latitude in the Netherlands (53°N) where both populations overlap. This is for *M. arvalis* the center (mid-latitude) of its distribution range, while our lab *M. oeconomus* are from a relict population at the lower boundary of its geographical range, which is an extension for this species to operate at southern limits. For this reason, local adaptation of the PNES may have evolved differently in the two species. The elevated *Tshr* expression and therefore the possible higher sensitivity to photoperiod in *M. oeconomus* raised under SP, might favour photoperiodic induction of reproduction earlier in the spring. This might be a strategy to cope with the extremely early spring onset at the low latitude for this relict *M. oeconomus* vole population.

Interestingly, the large peak in *Tshβ* expression (Fig. 2C) that is only observed in 21-day old LP *M. arvalis* males may be responsible for the drastic increase in testis weight when animals are 30 days old. Faster testis growth in LP *M. arvalis* males (Fig. 2C) might be induced by the 2-3 fold higher *Tshβ* levels compared to LP *M. oeconomus* males (Fig. 3C). However, this data have to be interpreted with caution since the current study only considered gene expression levels and did not investigate protein levels.

The reduced *Tshr* expression under LP early in life (Fig. 3E,F) may be induced by epigenetic mechanisms, such as increased levels of DNA methylation in the promoter of this gene, which will reduce its transcription. A role for epigenetic regulation of seasonal reproduction has been proposed based on studies of the adult hamster hypothalamus (Stevenson and Prendergast, 2013). We did not find any photoperiodic differences in overall hypothalamic DNA methyltransferase (i.e. *Dnmt1* and *Dnmt3a*) expression (Fig. S1G-J), which might be the result of sampling time. Previous studies in Siberian hamsters showed photoperiodic dependent circadian patterns of hypothalamic *Dnmt* expression (Stevenson, 2017). In order to study the effects of photoperiodic programming in development, DNA methylation patterns of specific promoter regions of photoperiodic genes at different circadian time points need to be studied in animals exposed to different environmental conditions earlier in development.

### Photoperiod induced changes in hypothalamic Dio2/Dio3 expression

The photoperiodic induced *Tshβ* and *Tshr* expression patterns are only reflected in the downstream *Dio2*/*Dio3* expression differences in the beginning of development (Fig. 3K,L), suggesting that this part of the pathway is sensitive to TSH at a very young age. However, *Dio2* and *Dio3* are also responsive to metabolic status, which can change as a consequence of changing DIO2/DIO3 levels. *M. oeconomus* and *M. arvalis* females show similar photoperiodic induced *Tshβ* patterns, while photoperiodic responses on *Tshr* are larger in *M. oeconomus*. The higher *Tshr* levels in *M. oeconomus* may be responsible for the higher *Dio2*, and lower *Dio3* levels in *M. oeconomus* females compared to *M. arvalis* females. However, the photoperiodic induced differences in gene expression levels between species is not reflected in female gonadal weight, indicating that additional signaling pathways are involved in regulating ovary and uterus growth. In males, *Dio2*/*Dio3* patterns are mainly determined by photoperiod, while different photoperiodic responses between species are lacking.

*Dio2* and *Tshβ* expression correlate only at the beginning of development (i.e. 15 and 21 days old) (Fig. 4A-D). These results are partly in agreement with the effects of constant photoperiod on hypothalamic gene expression in the Siberian hamster, showing induction of *Dio2* at birth when gestated under LP, and induction of *Dio3* at 15 days old when exposed to SP (Sáenz de Miera et al., 2017). Furthermore, it is thought that *Dio2/Dio3* expression profiles will shift due to both photoperiodic and metabolic changes rather than by constant conditions. Also, negative feedback on the Dio2/Dio3 system might be induced by changes in metabolic status. In wild populations of Brandt’s voles (*Lasiopodomys brandtii*), seasonal regulation of these genes, show elevated *Dio2/Dio3* ratios in spring under natural photoperiods, suggesting the crucial role for those genes in determining the onset of the breeding season in wild populations (Wang et al., 2019).

### Photoperiod induced changes in hypothalamic Kiss1 and Npvf expression

In females, both *Kiss1* and *Npvf* expression is higher under LP dependent on age (Fig. S1D,F), whereas in males no effects of photoperiod on these genes are found (Fig. S1C,E). Other studies report inconsistent photoperiodic/seasonal effects on ARC *Kiss1* expression in different species, which may be related to a negative sex steroid feedback on *Kiss1* expressing neurons (for review see, Simonneaux, 2020). For this reason, one might expect sex and species dependent levels of steroid negative feedback on both *Kiss1* and *Rfrp* expressing neurons in the caudal hypothalamus

In conclusion, our data show that somatic growth is photoperiodic sensitive in *M. oeconomus* while gonadal growth is photoperiodic sensitive in *M. arvalis. Tshr* expression is higher in the developing hypothalamus under SP conditions in both species, but more pronounced in *M. oeconomus*. Therefore, sensitivity to TSH might be programmed by photoperiod early in development. By programming TSH sensitivity early in life, animals would get the most appropriate reproductive response later in life. This expectation can be investigated by simulating spring and autumn changes in photoperiod in the laboratory and expose both vole species to these different photoperiod regimes. Both vole species program their PNES differently, dependent on photoperiod early in development, indicating that they use a different breeding strategy. *M. arvalis* uses a photoperiodic driven breeding strategy, while *M. oeconomus* might be using a more opportunistic breeding strategy.

## Acknowledgements

We would like to thank Saskia Helder for her valuable help in animal care.

## List of abbreviations

ARC: arcuate nucleus
*Dio2*: iodothyronine-deiodinase 2
*Dio3*: iodothyronine-deiodinase 3
*Dnmt1*: DNA methyltransferase 1
*Dnmt3a*: DNA methyltransferase 3a
GH: growth hormone
GnRH: gonadotropin-releasing hormone
*Kiss1*: Kisspeptin
KNDy: kisspeptin/neurokininB/Dynorphin
LP: long photoperiod
*Npvf*: neuropeptide VF precursor
PNES: photoperiodic neuroendocrine system
PT: pars tuberalis
SCN: suprachiasmatic nucleus
SP: short photoperiod
*Tshβ*: thyroid-stimulating-hormone-βsubunit

## References

Baker, J. (1938). The evolution of breeding seasons. Evol. Essays Asp. Evol. Biol. 161–177.

Dalum, J. Van, Melum, V. J., Wood, S. H. and Hazlerigg, D. G. (2020). Maternal photoperiodic programming: melatonin and seasonal synchronization before birth. Front. Endocrinol. (Lausanne). 10, 1–7.

Dardente, H., Wyse, C. A., Birnie, M. J., Dupré, S. M., Loudon, A. S. I., Lincoln, G. A. and Hazlerigg, D. G. (2010). A molecular switch for photoperiod responsiveness in mammals. Curr. Biol. 20, 2193–2198.

Dardente, H., Hazlerigg, D. G. and Ebling, F. J. P. (2014). Thyroid hormone and seasonal rhythmicity. Front. Endocrinol. (Lausanne). 5, 1–11.

Dardente, H., Wood, S., Ebling, F. and Sáenz de Miera, C. (2018). An integrative view of mammalian seasonal neuroendocrinology. J. Neuroendocrinol. 31,.

Dumbell, R. A., Scherbarth, F., Diedrich, V., Schmid, H. A., Steinlechner, S. and Barrett, P. (2015). Somatostatin Agonist Pasireotide Promotes a Physiological State Resembling Short-Day Acclimation in the Photoperiodic Male Siberian Hamster (Phodopus sungorus). J. Neuroendocrinol. 27, 588–599.

Ebling, F. J. P. (1994). Photoperiodic differences during development in the dwarf hamsters phodopus sungorus and phodopus campbelli. Gen. Comp. Endocrinol. 95, 475–482.

Evernden, L. N. and Fuller, W. A. (1972). Light alteration caused by snow and its importance to subnivean rodents. J. Zool. 50, 1023–1032.

Gall, C. Von, Stehle, J. H. and Weaver, D. R. (2002). Mammalian melatonin receptors: molecular biology and signal transduction. Cell Tissue Res 309, 151–162.

Gall, C. V. O. N., Weaver, D. R., Moek, J., Jilg, A., Stehle, J. H. and Korf, H.-W. (2005). Melatonin Plays a Crucial Role in the Regulation of Rhythmic Clock Gene Expression in the Mouse Pars Tuberalis. ann. N.Y. Acad. Sci. 511, 508–511.

Gerkema, M. P., Daan, S., Wilbrink, M., Hop, M. W. and Van Der Leest, F. (1993). Phase control of ultradian feeding rhythms in the common vole (Microtus arvalis): The roles of light and the circadian system. J. Biol. Rhythms 8, 151–171.

Hanon, E. A., Lincoln, G. A., Fustin, J.-M., Dardente, H., Masson-Pévet, M., Morgan, P. J. and Hazlerigg, D. G. (2008). Ancestral TSH mechanism signals summer in a photoperiodic mammal. Curr. Biol. 18, 1147–1152.

Hoffmann, K. (1973). Effects of short photoperiods on puberty, growth and moult in the Djungarian hamster (Phodopus sungorus). J. Reprod. Fertil. 54, 29–35.

Horton, T. H. (1984a). Growth and reproductive development of male Microtus montanus is affected by the prenatal photoperiod. Biol. Reprod. 31, 499–504.

Horton, T. H. (1984b). Growth and reproductive is affected development by the prenatal of male Microtus montanus photoperiod. Biol. Reprod. 504, 499–504.

Horton, T. H. (1985). Cross-fostering of Voles Demonstrates In Utero Effect of Photoperiod. Biol. Reprod. 33, 934–939.

Horton, T. H. and Stetson, M. H. (1992). Maternal transfer of photoperiodic information in rodents. Anim. Reprod. Sci. 30, 29–44.

Hut, R. A. (2011). Photoperiodism?: Shall EYA Compare Thee to a Summer’s Day? Curr. Biol. 21, R22–R25.

Hut, R. A., Paolucci, S., Dor, R., Kyriacou, C. P. and Daan, S. (2013). Latitudinal clines: an evolutionary view on biological rhythms. Proc. R. Soc. B Biol. Sci. 280, 20130433–20130433.

Korslund, L. (2006). Activity of root voles (Microtus oeconomus) under snow: Social encounters synchronize individual activity rhythms. Behav. Ecol. Sociobiol. 61, 255–263.

Król, E., Douglas, A., Dardente, H., Birnie, M. J., Vinne, V. van der, Eijer, W. G., Gerkema, M. P., Hazlerigg, D. G. and Hut, R. A. (2012). Strong pituitary and hypothalamic responses to photoperiod but not to 6-methoxy-2-benzoxazolinone in female common voles (Microtus arvalis). Gen. Comp. Endocrinol. 179, 289–295.

Malyshev, A. V, Henry, H. A. L. and Kreyling, J. (2014). Relative effects of temperature vs photoperiod on growth and cold acclimation of northern and southern ecotypes of the grass Arrhenatherum elatius. Environ. Exp. Bot. 106, 189–196.

Masumoto, K. H., Ukai-Tadenuma, M., Kasukawa, T., Nagano, M., Uno, K. D., Tsujino, K., Horikawa, K., Shigeyoshi, Y. and Ueda, H. R. (2010). Acute induction of Eya3 by late-night light stimulation triggers TSHβ expression in photoperiodism. Curr. Biol. 20, 2199–2206.

Nakane, Y. and Yoshimura, T. (2019). Photoperiodic Regulation of Reproduction in Vertebrates. Annu. Rev. Anim. Biosci. 7, 173–94.

Nakao, N., Ono, H., Yamamura, T., Anraku, T., Takagi, T., Higashi, K., Yasuo, S., Katou, Y., Kageyama, S., Uno, Y., et al. (2008). Thyrotrophin in the pars tuberalis triggers photoperiodic response. Nature 452, 317–322.

Ono, H., Hoshino, Y., Yasuo, S., Watanabe, M., Nakane, Y., Murai, A., Ebihara, S., Korf, H.-W. and Yoshimura, T. (2008). Involvement of thyrotropin in photoperiodic signal transduction in mice. Proc. Natl. Acad. Sci. 105, 18238–18242.

Peacock, J. M. (1976). Temperature and leaf growth in four grass species. J. Appl. Ecol. 13, 225–232.

Pfaffl, M. W. (2001). A new mathematical model for relative quantification in real-time RT-PCR. Nucleic Acids Res. 29, 16–21.

Phalen, A. N., Wexler, R., Cruickshank, J., Park, S. and Place, N. J. (2009). Photoperiod-induced differences in uterine growth in Phodopus sungorus are evident at an early age when serum estradiol and uterine estrogen receptor levels are not different. Comp. Biochem. Physiol. - A Mol. Integr. Physiol. 155, 115–121.

Prendergast, B. J., Gorman, M. R. and Zucker, I. (2000). Establishment and persistence of photoperiodic memory in hamsters. Proc. Natl. Acad. Sci. U. S. A. 97, 5586–5591.

Prendergast, B. J., Pyter, L. M., Kampf-Lassin, A., Patel, P. N. and Stevenson, T. J. (2013). Rapid induction of hypothalamic iodothyronine deiodinase expression by photoperiod and melatonin in juvenile Siberian hamsters (Phodopus sungorus). Endocrinology 154, 831–841.

Robson, M. J. (1967). A comparison of british and North African varieties of tall fescue (Festuca arundinacea). I. Leaf growth during winter and the effects on it of temperature and daylength. J. Appl. Ecol. 4, 475–484.

Sáenz de Miera, C., Bothorel, B., Jaeger, C., Simonneaux, V. and Hazlerigg, D. (2017). Maternal photoperiod programs hypothalamic thyroid status via the fetal pituitary gland. Proc. Natl. Acad. Sci. 114, 8408–8413.

Sáenz de Miera, C., Beymer, M., Routledge, K., Król, E., Selman, C., Hazlerigg, D. G. and Simonneaux, V. (2020). Photoperiodic regulation in a wild-derived mouse strain. J. Exp. Biol. 223, 1–9.

Sáenz De Miera, C. (2019). Maternal photoperiodic programming enlightens the internal regulation of thyroid-hormone deiodinases in tanycytes. J. Neuroendocrinol. 31, 12679.

Sáenz de Miera, C., Monecke, S., Bartzen-Sprauer, J., Laran-Chich, M. P., Pévet, P., Hazlerigg, D. G. and Simonneaux, V. (2014). A circannual clock drives expression of genes central for seasonal reproduction. Curr. Biol. 24, 1500–1506.

Scherbarth, F., Diedrich, V., Dumbell, R. A., Schmid, H. A., Steinlechner, S. and Barrett, P. (2015). Somatostatin receptor activation is involved in the control of daily torpor in a seasonal mammal. Am J Physiol Regul Integr Comp Physiol 309, 668–674.

Simonneaux, V. (2020). A Kiss to drive rhythms in reproduction. Eur. J. Neurosci. 51, 509–530.

Stetson, M. H., Elliott, J. A. and Goldman, B. D. (1986). Maternal transfer of photoperiodic information influences the photoperiodic response of prepubertal Djungarian hamsters (Phodopus sungorus sungorus). Biol. Reprod. 34, 664–669.

Stevenson, T. J. (2017). Circannual and circadian rhythms of hypothalamic DNA methyltransferase and histone deacetylase expression in male Siberian hamsters (Phodopus sungorus). Gen. Comp. Endocrinol. 243, 130–137.

Stevenson, T. J. and Prendergast, B. J. (2013). Reversible DNA methylation regulates seasonal photoperiodic time measurement. Proc. Natl. Acad. Sci. 110, 4644–4646.

Stumpfel, S., Bieber, C., Blanc, S., Ruf, T. and Giroud, S. (2017). Differences in growth rates and pre-hibernation body mass gain between early and late-born juvenile garden dormice. J. Comp. Physiol. B 187, 253–263.

Tkadlec, E. (2000). The effects of seasonality on variation in the length of breeding season in arvicoline rodents. Folia Zool. 49, 269–286.

van de Zande, L., van Apeldoorn, R. C., Blijdenstein, A. F., de Jong, D., van Delden, W. and Bijlsma, R. (2000). Microsatellite analysis of population structure and genetic differentiation within and between populations of the root vole, Microtus oeconomus in the Netherlands. Mol. Ecol. 9, 1651–1656.

Wang, D., Li, N., Tian, L., Ren, F., Li, Z., Chen, Y., Hu, X., Zhang, X., Song, Y., Hut, R. A., et al. (2019). Dynamic expressions of hypothalamic genes regulate seasonal breeding in a natural rodent population. Mol. Ecol.

Williams, L. M. and Morgan, P. J. (1988). Demonstration of melatonin-binding sites on the pars tuberalis of the rat. J. Endocrinol. 119, 1–3.

Wood, S. H., Christian, H. C., Miedzinska, K., Saer, B. R. C., Johnson, M., Paton, B., Yu, L., McNeilly, J., Davis, J. R. E., McNeilly, A. S., et al. (2015). Binary switching of calendar cells in the pituitary defines the phase of the circannual cycle in mammals. Curr. Biol. 25, 2651–2662.

Yellon, S. M. and Goldman, B. D. (1984). Photoperiod Control of Reproductive Development in the Male Djungarian Hamster (Phodopus sungorus)*. Endocrinology 114, 664–670.

Yellon, S. M. and Longo, L. D. (1987). Melatonin rhythms in fetal and maternal circulation during pregnancy in sheep. Am. Physiol. Soc.

